# Proteus software for physics-based protein design

**DOI:** 10.1101/2020.06.30.179549

**Authors:** David Mignon, Karen Druart, Vaitea Opuu, Savvas Polydorides, Francesco Villa, Thomas Gaillard, Eleni Michael, Georgios Archontis, Thomas Simonson

## Abstract

We describe methods and software for physics-based protein design. The folded state energy combines molecular mechanics with Generalized Born solvent. Sequence and conformation space are sampled with Replica Exchange Monte Carlo, assuming one or a few fixed protein backbone structures and discrete side chain rotamers. Whole protein design and enzyme design are presented as illustrations. Full redesign of three PDZ domains was done using a simple, empirical, unfolded state model. Designed sequences were very similar to natural ones. Enzyme redesign exploited a powerful, adaptive, importance sampling approach that allows the design to directly target substrate binding, reaction rate, catalytic efficiency, or the specificity of these properties. Redesign of tyrosyl-tRNA synthetase stereospecificity is reported as an example.

## 1 Introduction

Computational protein design (CPD) is an exciting field that has had spectacular successes. Whole proteins have been redesigned, ^1–6^ new enzymes obtained,^7,8^ a new fold produced, ^2^ designed enzymes that digest plastic have reached industrial processes. ^9^ However, failures are also common. For example, the success rate for small protein redesign in a recent, large-scale study was about 6%.^6^ Redesigned enzymes often have low activities.^10,11^ Thus, progress is still very much needed. Many applications have used a large dose of empiricism, by including terms in the energy function that are statistical, or knowledge-based. Some were derived from amino acid propensities or pair propensities found in sequence alignments;^12^ others were derived from structural propensities in X-ray structures. For example, one recent energy function included terms derived from torsion angle and hydrogen-bond distance distributions in crystal structures, ^5,13^ with the logarithm of the experimental probability distribution treated as an effective energy.

A complementary route is to put more emphasis on physics-based ingredients. This includes the energy function, but also the strategy to sample sequences and conformations. If one uses Monte Carlo, sampling can be made to follow a Boltzmann distribution, which has two consequences. First, powerful importance sampling techniques can be used, such as Wang-Landau adaptive MC.^14^ Second, sequences can be sampled or ranked according to relative free energies, such as binding or folding free energies. This approach, which combines physics-based energy with Boltzmann sampling, we refer to as “physics-based” protein design, or *ϕ*-CPD. While many of the ingredients are old, others were developed recently. Their advantage is that they provide good accuracy, are transferable to all biomolecules, can be systematically improved, and give physical insights.

Here, we present the Proteus software for *ϕ*-CPD (available from https://proteus.polytechnique.fr) and associated methodology. We first recall some theoretical background. We begin with the physical interpretation of a MC simulation in sequence space. We recall the model for the folded state, which uses a fixed backbone, discrete side chain rotamers, a molecular mechanics protein energy, and one of two main implicit solvent models. We recall the unfolded state model, including a maximum likelihood parameterization framework. We describe two advanced MC variants: hybrid MC for multi-backbone design and adaptive MC sampling for the design of protein–ligand binding.

Next, we briefly decribe the software itself. Basic information is in an earlier article.^15^ A major new release (3.0) was finished in 2019,^16^ with several new functionalities, including an improved solvent model^17,18^ and advanced sampling methods, partly developed by our groups. Thanks to all these, we recently reported the first successful whole protein redesign with a physics-based energy function. ^19^

Finally, we present examples of two important CPD applications. One is the complete redesign of a small set of PDZ proteins, using the two main variants of the solvent model. The other is the redesign of the tyrosyl-tRNA synthetase to maximize its activity for a D-tyrosine substrate, instead of the usual L-tyrosine. Thanks to the Wang-Landau adaptive MC and a structural model of the transition state, designed sequences are populated according to their catalytic efficiency, with efficient variants exponentially enriched.

## 2 Theoretical methods

### 2.1 What is sequence space?

Sequence design with Proteus is done by running long MC simulations where selected amino acid positions can mutate freely. Before any practical details, we consider the physical meaning of an MC exploration of sequence space and its statistical ensemble. In the simulation, one copy of the folded protein is explicitly represented. The unfolded state is included implicitly, with a representation that does not involve a 3D structural model (see below). The simulation is propogated with the energy function *E_M_* = *E_f_* − *E_u_* (the folding energy). One possible move is a mutation: we modify the sidechain type 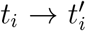 at a chosen position *i* in the folded protein, assigning a particular rotamer 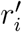 to the new sidechain. The energy change is

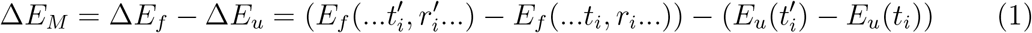

Δ*E_M_* measures the stability change due to the mutation (for the given set of rotamers); it is as if we performed the reverse mutation 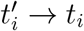 in the unfolded form. For two sequences *S*, *S*′, the probability ratio over a long simulation is

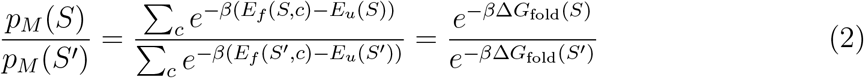

where the sums are over all conformations *c*. We recognize the ratio of Boltzmann factors for *S* and *S*′ folding. Δ*G*_fold_(*S*) denotes the folding free energy of sequence *S* (respectively, *S*′). Eq. (2) has a simple interpretation: the simulation, with the energy function *E_M_* = *E_f_* − *E_u_* and appropriate move probabilities, leads to the same distribution of states as a macroscopic, equilibrium, physical system where all sequences *S*, *S*′, … are present at equal concentrations, and are distributed between their folded and unfolded states according to their relative stabilities. This is precisely the situation we want to mimic. In this interpretation, a Monte Carlo mutation move *S* → *S*′ amounts to unfolding one copy of *S* and refolding one copy of *S*′ (Fig. 1). Thus, MC in “sequence space” actually takes place in a conformational space, where ordinary statistical mechanics apply.

**Figure 1:**
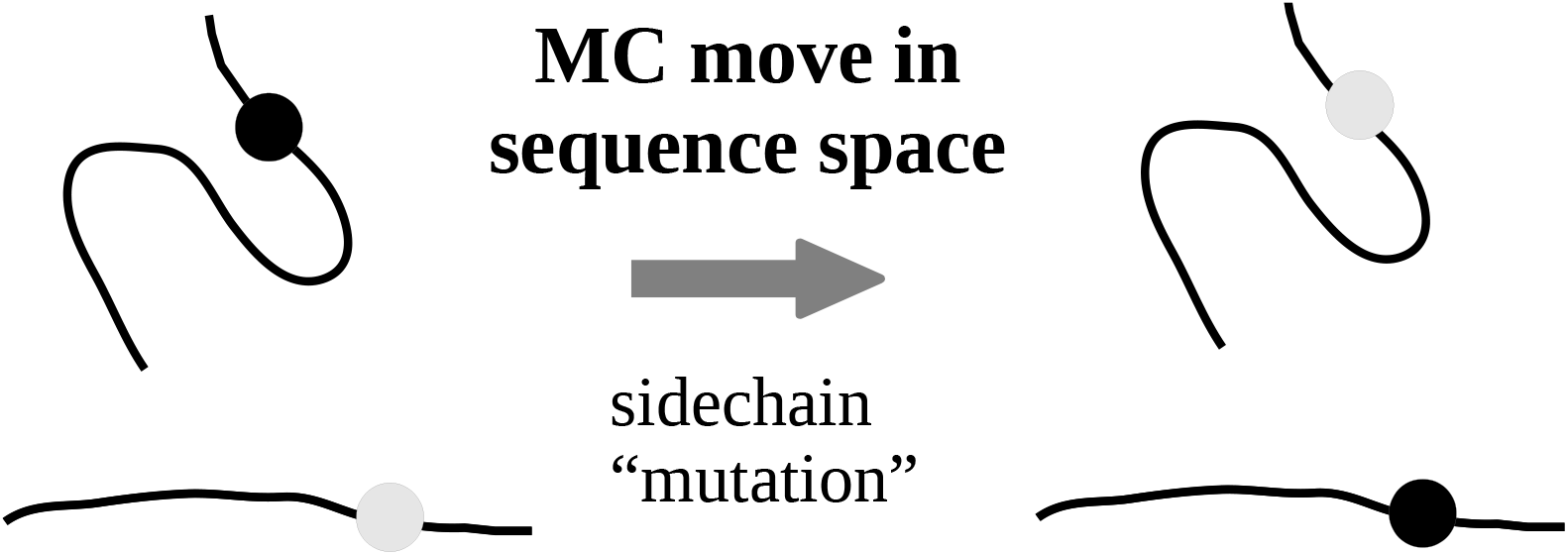
A MC mutation move: a point mutation is performed in the folded state, along with the inverse mutation in the unfolded state.

### 2.2 Structure and energy model for the folded state

Proteus uses a model with one or a few fixed backbone conformations and a discrete rotamer library that can include native rotamers (the X-ray conformation of a side chain). Only the Tuffery library^20,21^ is currently provided with Proteus; others can be used at the cost of some reformatting. Design with a fully flexible backbone and continuous rotamers was implemented in Proteus recently, ^22^ using a hybrid MC/MD approach. However, the current implementation is still slow and is not described here.

The energy function for the folded state has the form *E*_folded_ = *E*_intra_ + *E*_solv_. The first term is the protein internal energy from the Amber ff99SB force field^23^ (slightly modified for CPD^24^). The second term is the contribution of solvent, *E*_solv_ = *E*_GB_ + *E*_nonpolar_. The first, Generalized Born (GB) term captures the main electrostatic effects:^25,26^

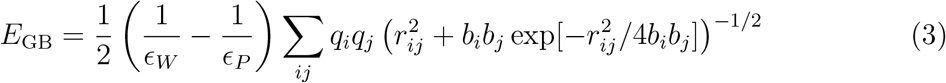

where *ϵ_W_* and *ϵ_P_* are the solvent and protein dielectric constants; *r_ij_* is the distance between atoms *i, j* and *q_i_*, *b_i_* are the charge and “solvation radius” of atom *i*.^25,26^ The second solvent term *E*_nonpolar_ represents dispersion and hydrophobic effects. It can be a surface area (SA) term:

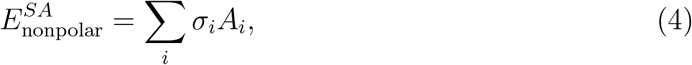

where *A_i_* is the solvent accessible surface area of atom *i*; *σ_i_* is a parameter that reflects each atom’s preference to be exposed. A Lazaridis-Karplus (LK) form is also possible:^18,27^

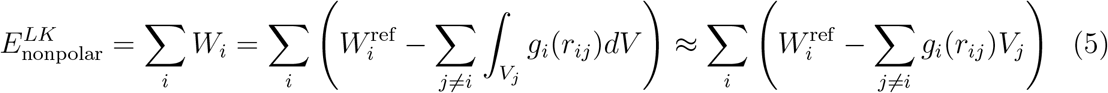

where the sum is over all atoms *i*, 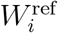 is an empirical value for a fully exposed atom *i*, *g_i_*(*r_ij_*) is a gaussian energy density that decays exponentially with the separation *r_ij_*, and *V_j_* is the volume of atom *j*,

Both the GB and SA terms have a many-body character. In the GB term, the solvation radius *b_i_* approximates the distance from *i* to the protein surface and is a function of the coordinates of all the protein atoms. In the SA term, the same area on one atom can be buried by two others. For SA, Proteus provides either a standard approximation by a pairwise additive form^1,26^ or a less-standard, slightly more accurate approximation.^24^ For GB, Proteus provides a pairwise approximation, called NEA (“Native Environment Approximation”) where the solvation radius *b_i_* is computed ahead of time, with the rest of the system in its native sequence and conformation. An exact method is also provided, called FDB (“Fluctuating Dielectric Boundary”), where the *b_i_* are computed on-the-fly, during MC, with no approximation.

With a pairwise additive energy, a fixed backbone and a discrete rotamer library, residue interaction energies can be computed ahead of time and stored in a lookup table, or “energy matrix”.^1^ This makes subsequent MC exploration very efficient. For the many-body GB variant FDB, an energy matrix can still be used, with the GB energies represented by a lookup table of lookup tables.^17,28^

With the discrete rotamer picture, side chain pairs only have a few available arrangements. For nearby pairs, this can lead to steric clashes, which would be resolved by departing slightly from the rotamers. Most CPD methods address this either by modifying the form of Lennard-Jones interactions at short range^13^ or by reducing the van der Waals radius of atoms. Proteus uses a different method:^29^ for each pair of residues *I*, *J* and each rotamer combination, a short energy minimization is performed (around 15 conjugate gradient steps), with the backbone fixed, including only the interactions within and between the pair.^15,24^ The *I* − *J* interaction energy is computed after the minimization and stored in the energy matrix.

### 2.3 The unfolded state

#### 2.3.1 Extended peptide model

For the unfolded state, we use a model that is widespread, where each amino acid interacts with solvent but not with the other amino acids. This is roughly the situation with an extended peptide, where each side chain is exposed to solvent but distant from other side chains and from backbone groups other than its immediate neighbors. For a sequence *S*, the unfolded state energy then has the form:

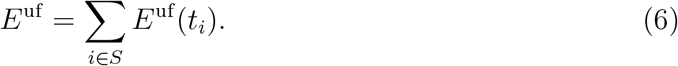

The sum is over all amino acids; *t_i_* represents the sidechain type at position *i*. The type-dependent quantities 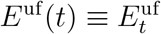 are sometimes referred to as “reference energies”. They can be computed from the 3D structure of an extended peptide. Such an approach is sufficient for the design of protein–ligand binding.^30^

#### 2.3.2 Knowledge-based model

For whole protein design, the folding energy is a key quantity and the extended peptide unfolded model is not accurate enough. Instead, the 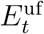 can be chosen empirically, to reproduce the amino acid composition of a set of experimental proteins. This was done in most successful whole protein designs, including those with Proteus. ^19^ The 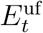 can be thought of as effective chemical potentials of each amino acid type. The MC method employed by Proteus generates a Markov chain of states^31,32^ such that the folding energies follow a Boltzmann distribution. This makes it possible to choose the reference energies that maximize the probability of the experimental sequences.

Let 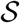 be a set of *N* experimental, “target” sequences *S*. The Boltzmann probability of *S* is

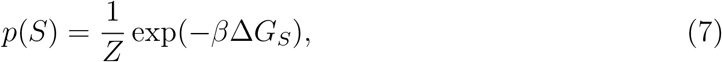

where 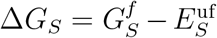 is the folding free energy of *S*, 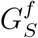 is the free energy of the folded form, *β* = 1*/kT* is the inverse temperature and *Z* is a normalizing constant (the partition function). We denote 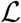 the probability of the entire sequence set, which depends on the model parameters 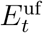; we refer to 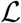 as their likelihood.^33^ A necessary condition to maximize 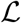 is that its derivatives with respect to the 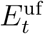 should be zero. We obtain^34^

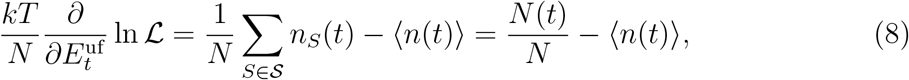

where *n_S_*(*t*) is the number of amino acids of type *t* within the sequence *S* and *N* (*t*) is the number in the whole dataset 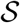. Therefore,

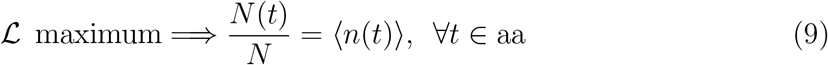

To maximize 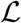, we should choose 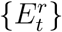 such that a long simulation gives the same amino acid frequencies as the target database. Various gradient methods can be used to search for the best 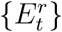; one is detailed in Results. Second derivatives are also readily obtained:

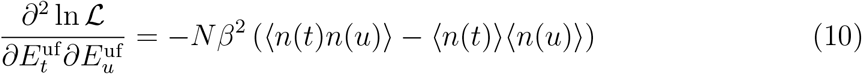

Thus, the 2nd derivatives of the log likelihood with respect to the 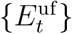 are given by the covariances of the conjugate quantities {*n_S_*(*t*)}, as usual in statistical physics.

### 2.4 MC and REMC simulations

Proteus simulations use one- and two-position moves, where either rotamers, amino acid types, or both are changed. The mutating positions are user-defined and depend on the specific problem. Sampling can be enhanced by Replica Exchange Monte Carlo (REMC), where several MC simulations are run in parallel, at different temperatures. ^35^ Periodic swaps are attempted between the conformations of two replicas (adjacent in temperature) and accepted or rejected by a Metropolis test. REMC uses OpenMP parallelization. Thanks to precalculation of the energy matrix, MC with Proteus is very fast, and a billion steps can be run in a few hours on a desktop machine.

Boltzmann sampling depends on the detailed balance property:^36^ in a long simulation, the number of transitions between any two states should be the same in each direction. This is guaranteed^37^ if the move set allows any state to be reached (eventually) from any other (as here), if the system is “aperiodic” (there are no states that form “periodic orbits,” trapping the system indefinitely), and if the Kolmogorov reversibility condition is verified: the products of probabilities around closed loops of states are the same in both directions (as here).^35^ Aperiodicity is unproven but always verified in our experience.

### 2.5 Hybrid MC for multi-backbone design

With fixed-backbone *ϕ*-CPD, backbone deformation is not neglected, but treated implicitly through the protein dielectric constant. This treatment is not sufficient for some problems; for example, it does not allow the backbone to shift and make room for large new side chains. One way to introduce backbone deformation explicitly is to perform simulations where the backbone conformation can vary within a small set of possibilities. This is known as multi-backbone design. To sample sequence, rotamer, and backbone degrees of freedom all at once is a hard problem. Proteus provides a hybrid MC method, with backbone moves in addition to rotamer and mutation moves.^38^ A hop to a new backbone conformation will rarely be accepted, since the new backbone will usually require side chain rotamer adjustments. Therefore, Proteus uses a complex, two-stage move, where a hop to a new backbone is followed by a short MC segment (50–100 steps) where rotamers (and possibly sequence) evolve and relax. Only at the end is an acceptance test performed, which accepts or rejects the entire move. The acceptance probability takes the form of a path integral, which is computed using a simple approximation ^39^ or a more complex one.^38^ Both involve estimating the probability of selecting the relaxed conformation as a trial state. The first approximation simply multiplies the probabilities of the elementary MC steps that were taken along the relaxation segment. The second averages over an ensemble of ∼100 possible relaxation segments. The quality of the approximation can be improved systematically by increasing the length of the relaxation segment. Numerical tests on small proteins showed that Boltzmann sampling was achieved accurately with a relaxation length of around 100 steps.^38^

### 2.6 Adaptive Wang-Landau MC for protein-ligand binding

Protein-ligand binding is a main application of CPD. Most studies have used a simple approach where the energy of the complex is targeted for optimization. However, the property to optimize is quite different: the binding free energy, which involves a competition between bound and unbound states. An efficient and elegant approach was proposed recently.^30,40^ Using an adaptive, Wang-Landau MC procedure^14^ to “learn” an appropriate bias, one can rapidly obtain large sequence ensembles populated according to a ligand binding free energy. The bias *B* is constructed gradually over a simulation of the unbound protein, such that all sequences reach comparable populations. *B* is then essentially the sequence free energy with its sign changed. Next, the bias is included in a simulation of the complex, where it “subtracts out” the unbound state. Thus, positive design of the bound state and negative design of the unbound state are achieved. Remarkably, the result is a Boltzmann distribution where sequence populations measure their affinities. The method can be readily extended to design for ligand binding specificity. In particular, one can design for specific binding of activated ligands that form the transition state of an enzyme reaction. Thus, one can design directly for catalytic power. The first application was reported very recently.^41^ Another is given in Results, where the tyrosyl-tRNA synthetase enzyme is redesigned to maximize activity towards the D-tyrosine substrate, instead of the usual L-tyrosine.

In practice, we perform a long MC simulation of the apo protein, where a few positions can mutate, arbitrarily numbered 1, …, *p*. We gradually increment a bias potential until all the side chain types at the mutating positions have roughly equal populations, thus flattening the free energy landscape. The bias potential contains terms associated with each mutating position and possibly pairs of positions. ^41^ The individual terms are updated at regular intervals of length *T*. At each update, whichever sequence variant (*s*_1_(*t*), *s*_2_(*t*), …, *s_p_*(*t*)) is populated is penalized by adding an increment to each corresponding term in the bias. The increments decrease exponentially as the bias increases, as in well-tempered metadynamics.^42,43^ Over time, the bias for the most probable states grows until it pushes the system into other regions of sequence space.

In a second stage, we simulate the protein-ligand complex in the presence of the bias. The binding free energy relative to a reference sequence *S_r_*, in the presence of the bias, has the form:

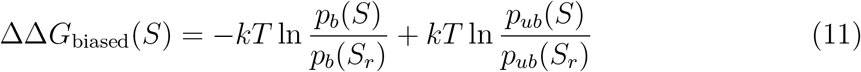

The relative binding free energy in the *absence of bias* has the form:

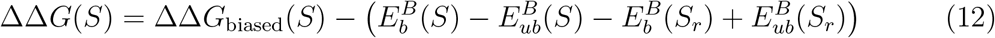

Subscripts *b*, *ub*, indicate the bound or unbound state, respectively. Since the bias is (usually) the same in both states, it cancels out from ΔΔ*G*_biased_(*S*), which is equal to the relative binding free energy *in the absence of bias*, ΔΔ*G*(*S*). While the bias may disappear from (11), it is essential for accurate sampling. Perfect flattening, however, is not usually achieved, nor is it needed.

## 3 Overview of the software implementation

Proteus is available from https://proteus.polytechnique.fr, with source code, binaries, and a detailed manual. A series of tutorials are available, with input and output. One covers redesign of a small peptide, where a few positions are allowed to mutate freely, with folding energy as the design target. One covers the full redesign of a PDZ domain, using previously determined reference energies. One covers the redesign of an enzyme for substrate binding. It can easily be adapted to select for transition state binding or binding specificity. One covers acid/base calculations for the BPTI protein. Finally, the most complex covers the optimization of unfolded energies for whole protein redesign. Tutorials for multi-backbone design are not yet available.

Proteus has four components:

1. the molecular simulation program **protX**, mostly written in Fortran 90; controlled by its own script language, borrowed from X-PLOR;^44^
2. a set of scripts in the protX scripting language that control the calculation of an energy matrix for the system of interest;^1^
3. a C program, **protMC** for exploring the space of sequences and conformations using various search algorithms, including Monte Carlo (MC); controlled by an xml script;
4. a collection of perl, python, and shell scripts that automate various steps.

The flowchart for a design calculation is schematized in Fig. 2. More details can be found in previous articles^15^ and the Proteus manual. ^16^ From the protein 3D structure (PDB file), the sequence is read and the topology or 2D “structure” is constructed and stored in a “protein structure file” or PSF, equivalent to the one used by Charmm, X-PLOR, and NAMD.^44–46^ Positions that will be allowed to mutate are indicated in a protX stream file sele.str. At these positions, “giant” residues are constructed, which carry multiple side chains. Intermediate files are stored, then the diagonal and off-diagonal elements of the energy matrix are computed by the protX scripts matrixI.inp and matrixIJ.inp. ProtMC is run to produce an MC trajectory, which has the form of a time series of rotamer lists. Post-processing is done, mainly by perl and python scripts. 3D structures corresponding to selected MC snapshots can be constructed with protX for analysis. More details can be found in the documentation. ^16^ Several tests and benchmarks were reported recently, including side chain reconstruction,^21^ acid/base calculations, ^17^ and successful whole protein redesign.^19,35,47^ Performance was comparable to several well-established tools.

**Figure 2:**
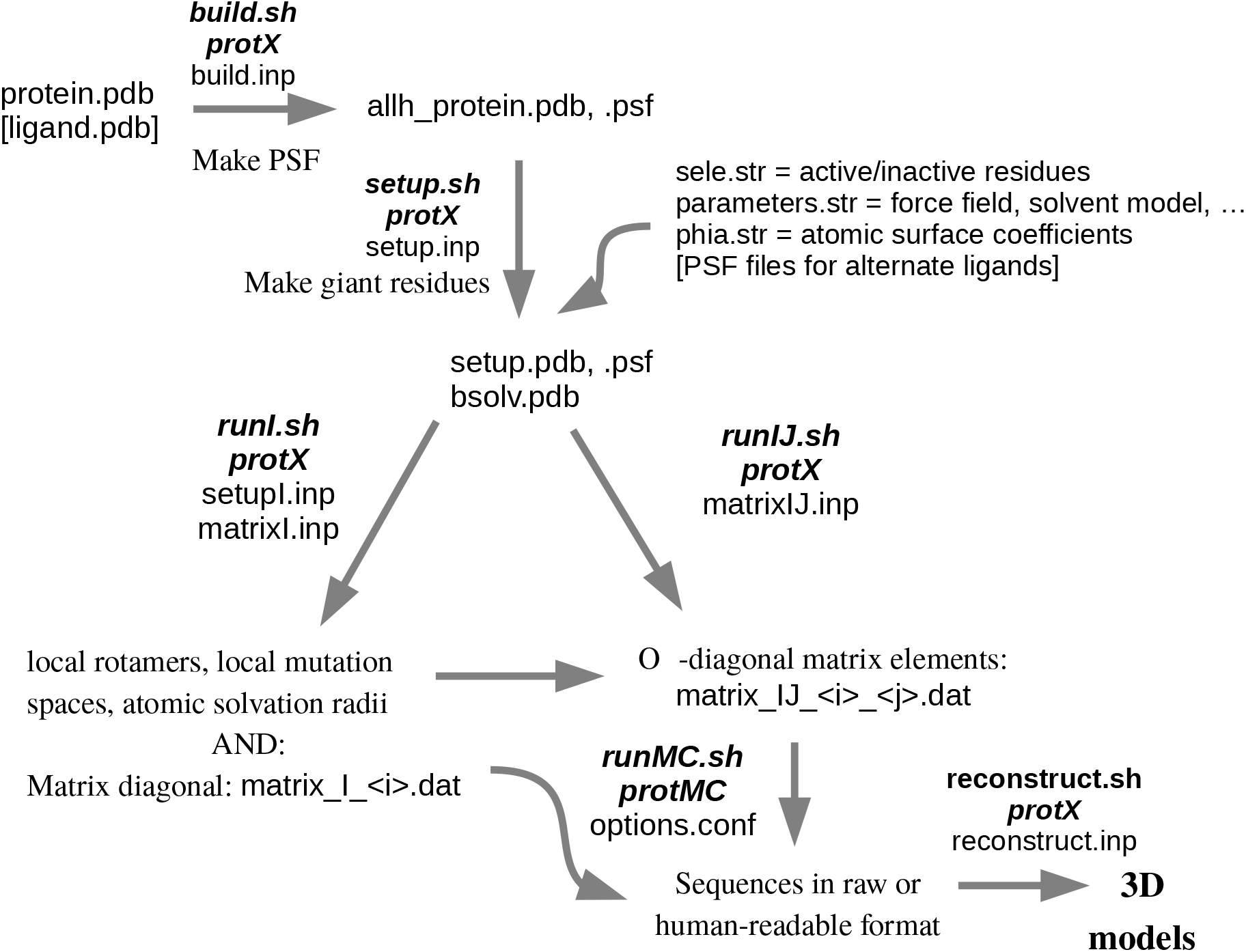
Flow chart for the energy matrix and sequence generation.

## 4 Whole protein redesign

We describe briefly a new set of whole protein design benchmarks, which illustrate the optimization of empirical unfolded energies and give an idea of the main options for implicit solvent treatment. The test proteins are three PDZ domains (Table 2). The unfolded energies are optimized specifically for this family.

### 4.1 Unfolded model and solvent treatment

The unfolded state plays a key role because whenever a mutation occurs in the folded structure, the reverse mutation occurs in the unfolded structure (Fig. 1). The unfolded energy is a sum of independent contributions from all residues, which depend on the side chain type but not the 3D structure. Let 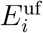 be the contribution of residue *i*. This contribution depends on the side chain type *t_i_* at position 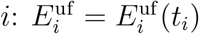. Often, we assume there is no dependency on the residue number: 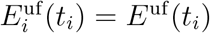. With this assumption, the unfolded state is characterized by a set of “unfolded energies” *E*^uf^ (*t*) that depend on the side chain type *t* but not its position in the polypeptide chain. The values *E*^uf^ (*t*) are normally chosen so that a simulation will reproduce the overall amino acid composition of a set of natural homologs, maximizing their likelihood. In the present benchmarks, we apply an unfolded model where amino acid positions are grouped according to their buried or exposed character in the folded protein. Each group will have its own set of type-dependent unfolded energies. This model assumes that the folded structure affects the unfolded model, either because residual structure is retained in the unfolded state, or because the folded model compensates for some of the errors in the unfolded model.^34^

The idea is to start from an initial set of unfolded energies, run a set of 2–3 MC simulations for each protein and compare the computed composition to the target one. The unfolded energies are then updated, by adding an increment that is related to the likelihood gradient. The procedure is repeated: a new set of MC simulations is done, the composition computed, and the unfolded energies updated. The procedure is run until convergence. The *E*^uf^ (*t*) update rule was the “linear” rule:^34^

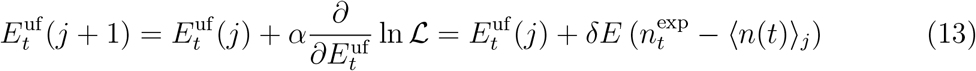

Here, *j* is an iteration number; *α* is a constant; 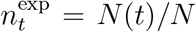 is the mean population of amino acid type *t* in the natural sequences; ⟨⟩_*j*_ indicates an average over a simulation done using the current unfolded energies 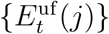, and *δE* is an empirical constant with the dimension of an energy, referred to as the update amplitude. We have omitted the distinction between buried and exposed positions here for simplicity. In each MC simulation, every other amino acid position was active, while the rest were inactive. Thus, there were two sets of active positions (per protein), and one simulation was done for each set. The overall computed amino acid composition was thus obtained at each iteration from six simulations, two per protein. In early iterations, amino acid types were grouped into 11 classes, with one adjustable unfolded energy per class, and a large *δE* was used. In later iterations, each amino acid type had its own adjustable unfolded energy and a smaller *δE* was used. Details are in Supplementary Material.

We compared several implicit solvent treatments for the folded state. They all included GB electrostatics, with different protein dielectric values *ϵ_P_*. One used the Lazaridis-Karplus nonpolar term (LK). The others used a surface area nonpolar term (SA) with an exact SA calculation. The LCPO approximation^48^ found in Amber^49^ gaven-poorer results that are not reported. With the exact SA, triple overlaps involving two side chains and backbone were either computed explicitly ^24^ (the Proteus default) or ignored (pairwise approximation, or SApw), which is more efficient.

### 4.2 Whole protein design results

The unfolded energies obtained with the different solvent treatments are given in Supplementary Material. Once they were optimized, the three PDZ proteins were redesigned by running a long MC simulation where all positions (except Gly, Pro) could mutate into all types (except Gly, Pro). To characterize the designs, we focus on the top 10000 sequences (the largest folding energies) for each protein. The mean amino acid composition obtained is in Supplementary Material. Similarities to the Pfam database^50^ are shown in Table 1 and Fig. 3A. Scores obtained with the Superfamily fold recognition tool are in Table 2. Sequence logos are in Fig. 3B. Mean sequence entropies are in Table 3. They use six residue classes, instead of the usual 20 types (see Supplementary Material).

**Table 1:** Mean similarity scores compared to Pfam.

|  | NHERF |  | Syntenin |  | DLG2 |  |
| --- | --- | --- | --- | --- | --- | --- |
|  | All | Core | All | Core | All | Core |
| SA<sup>a</sup> | 9.6 | 35.1 | 14.5 | 32.3 | -2.6 | 32.6 |
| SApw | 14.5 | 38.5 | 9.7 | 34.4 | -7.0 | 32.3 |
| SA | 16.0 | 36.3 | 14.3 | 32.1 | -5.6 | 31.6 |
| LK | 21.0 | 39.7 | 15.5 | 37.4 | 4.2 | 43.0 |
| Pfam | 47.2 | 31.6 | 60.9 | 31.6 | 39.8 | 31.6 |

Mean Blosum40 similarity between designed sequences and the PDZ Pfam alignment, averaged over all or core positions. <sup>a</sup>Used $\epsilon_P=4$ .

**Table 2:** Designed sequence Superfamily scores.

| PDZ<br>protein | PDB<br>code | Nonpolar<br>model | Match<br>length<sup>a</sup> | Superfamily<br>E-value<sup>b</sup> | Family<br>E-value<sup>b</sup> | Success<br>rate<sup>c</sup> |
| --- | --- | --- | --- | --- | --- | --- |
| NHERF | 1G9O | SA<sup>d</sup> | 77/91 | $1 \times 10^{-9}$ | $4 \times 10^{-3}$ | 100% |
| Syntenin | 1R6J | SA<sup>d</sup> | 72/82 | $3 \times 10^{-7}$ | $5 \times 10^{-3}$ | 82% |
| DLG2 | 2BYG | SA<sup>d</sup> | 81/97 | $1 \times 10^{-4}$ | $4 \times 10^{-3}$ | 80% |
| NHERF | 1G9O | SAPw | 79/91 | $8 \times 10^{-13}$ | $5 \times 10^{-3}$ | 100% |
| Syntenin | 1R6J | SAPw | 74/82 | $3 \times 10^{-3}$ | $8 \times 10^{-3}$ | 100% |
| DLG2 | 2BYG | SAPw | 59/97 | $1 \times 10^{-6}$ | $3 \times 10^{-3}$ | 100% |
| NHERF | 1G9O | SA | 83/91 | $2 \times 10^{-12}$ | $5 \times 10^{-3}$ | 100% |
| Syntenin | 1R6J | SA | 70/82 | $7 \times 10^{-7}$ | $5 \times 10^{-3}$ | 100% |
| DLG2 | 2BYG | SA | 78/97 | $2 \times 10^{-8}$ | $4 \times 10^{-3}$ | 100% |
| NHERF | 1G9O | LK | 82/91 | $1 \times 10^{-17}$ | $8 \times 10^{-2}$ | 100% |
| Syntenin | 1R6J | LK | 75/82 | $1 \times 10^{-11}$ | $4 \times 10^{-3}$ | 100% |
| DLG2 | 2BYG | LK | 90/97 | $6 \times 10^{-15}$ | $2 \times 10^{-2}$ | 100% |

<sup>a</sup>The average match length for sequences recognized by Superfamily and the total sequence length. <sup>b</sup>Average E-values for Superfamily assignments to the correct SCOP superfamily/family. <sup>c</sup>The number of designed sequences (out of 10000 tested) assigned to the correct SCOP superfamily/family. <sup>d</sup>Using the pairwise approximation.

**Table 3:**
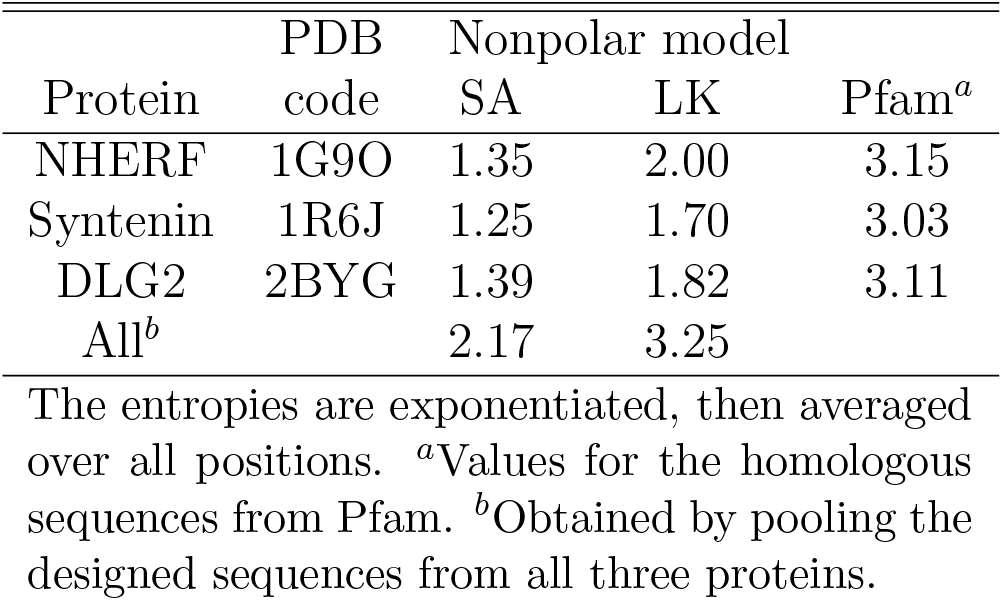
Designed and Pfam sequence entropies.

**Figure 3:**
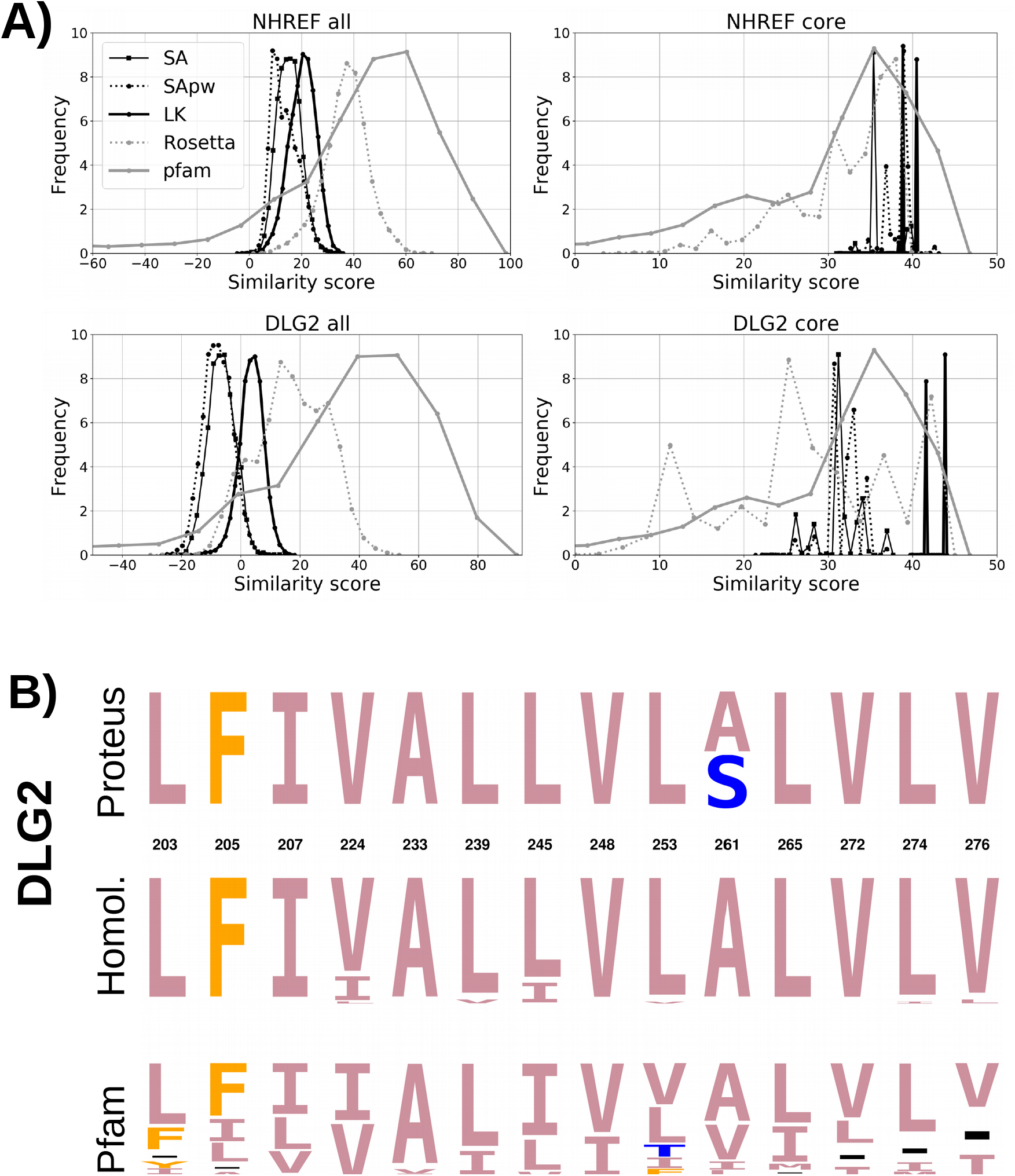
**A)** Histograms of Blosum40 similarity scores to Pfam sequences for two designed PDZ proteins: NHERF and DLG2. The similarity between Pfam sequences is also shown (RP-55 Pfam alignment). Design used either GBSA or GBLK. **B)** Sequence logos for the core region of DLG2. CPD sequences (GBLK), close homologs and full Pfam set.

The best performance was obtained with GBLK, followed by GBSA(*ϵ_P_* =8), with or without the pairwise approximation. Notice that the LK energy term was parameterized earlier^18^ using structural and thermodynamic data, not CPD data. With both the GBSA(*ϵ_P_* =8) and GBLK models, all the designed sequences were classified as PDZ domains by Superfamily, and the Blosum similarity to Pfam sequences for the protein cores was very high—higher even than the Pfam-to-Pfam similarity. For the whole protein, similarity to Pfam was lower, but still within the range of scores seen for some experimental sequences (Fig. 3A). Values obtained earlier^17^ with Rosetta^51^ are also shown. The GBLK similarities to Pfam are lower than Rosetta overall, but higher for the core. Sequence logos for one example (DLG2; Fig. 3B) show perfect recovery of the native core. Sequence entropies (Table 3) were lower than within the collection of natural sequences. However, GBLK gave a higher diversity than GBSA, and when the sequences designed with GBLK using all three backbones were pooled, the entropy was close to that of the Pfam set. Thus, with three PDZ backbones, GBLK design recapitulated, on average, the natural diversity.

## 5 Enzyme redesign: the stereospecificity of tyrosyl-tRNA synthetase

Tyrosyl-tRNA synthetase (TyrRS) catalyzes the attachment of tyrosine (Tyr) to its cognate tRNA^Tyr^, which carries the appropriate triplet of nucleotides as its anticodon, helping establish the genetic code. The attachment occurs through two successive reactions, which can occur separately. Here, we only consider the first, which forms tyrosyl adenylate (TyrAMP) from Tyr and ATP. TyrRS has been extensively engineered to accept unnatural amino acids as substrates, leading to genetic code expansion. ^52,53^ Earlier, we used Proteus to redesign its sterospecificity, and discovered a point mutant that prefers D-Tyr over the usual L-Tyr as its substrate. ^10^ That study used as a design criterion the energy of the Tyr:TyrRS complex. Although it successfully produced inverted stereospecificity, the redesigned enzyme had a weak activity, like many other redesigned enzymes. ^7,8,11^ This may be due to the use of the total energy as the design criterion.

Now, thanks to the new adaptive MC procedure, ^41^ we can use more relevant quantities. We can directly target the substrate affinity, the catalytic rate, or the catalytic efficiency, which is the ratio of the rate and the Michaelis constant, and closely approximates the second order rate constant. ^54^ We can also target the specificity ratio of the D-Tyr and L-Tyr values of any of these quantities. In each case, an adaptive bias is used to flatten the free energy landscape of a “reference state”, such as the apo enzyme. A “target state”, such as the transition state complex is then simulated, including the bias. Designs that have the most favorable transition state binding are then preferably sampled. When designs are selected for one property, it often happens that other properties like protein stability, may be degraded. This can easily be overcome by filtering designs *a posteriori*, and discarding those that are predicted to be unstable. Notice that a non-empirical, extended peptide, unfolded model is sufficient for this filtering step. The knowledge-based model is not needed, and in fact, the unfolded, or reference energies do not play any role in the MC simulations. (They can be set to zero, although the bias converges more rapidly if more physical values are used.)

We applied this strategy to *E. coli* TyrRS. Four active site residues were allowed to mutate freely, as previously:^10^ Asp81, Tyr175, Gln179, Gln201. Several design criteria were used (Table 4). In each case, the landscape of a suitable reference state was flattened. The bias energies were then used in the design of a target state:

**Design for affinity:** The free energy landscape of the apo enzyme (reference state) was flattened by an adaptive bias. Then the L-Tyr or D-Tyr complex was simulated (target state), including the bias. Since the bias approximates the negative free energy of the apo state, tight-binding variants were exponentially enriched. Sampled variants with low stability were discarded *a posteriori* (filtering step). Specifically, we discarded sampled sequences whose folding energy (estimated with an extended peptide unfolded model) was degraded, relative to WT.
**Design for catalytic efficiency:** As reference state, we used the apo enzyme (no ATP or Tyr). The landscape was flattened adaptively as before. Then, the L-Tyr or D-Tyr transition state complex (target state) was simulated. We actually flattened the landscape of the target state as well (with its own adaptive bias; see Methods in Supplementary Material). This allowed us to sample as many variants as possible for the target state (not only the top few). Sampled variants with low stability were filtered out *a posteriori*.
**Design for specificity:** The target property is the ratio of *k*_cat_*/K_M_* values for D-Tyr and L-Tyr. The activated L-Tyr complex (reference state) was flattened; then the activated D-Tyr complex (target state) was simulated, including the bias. Selection for specificity can yield variants that have good specificity (D-Tyr preference) but low D-Tyr activity. Therefore, we finished by filtering out and discarding variants with low stability *or* low D-Tyr activity or efficiency (lower than the WT).
**Design for specificity (more aggressive):** A more complex and aggressive protocol is possible, with three, not two simulations. First, we perform catalytic efficiency design separately for both L- and D-Tyr, making sure to flatten the apo and both transition states (three simulations). Next, we rank the sampled variants based on specificity. Because all states were flattened and aggressively sampled, we expect to visit not only the most specific variants, but also the most active. We finish by filtering out variants with low stability or D-Tyr activity.

**Table 4:** TyrRS design protocols.

| Design criterion | Binding |  | Rate |  | Efficiency |  | Specificity |  |
| --- | --- | --- | --- | --- | --- | --- | --- | --- |
|  | L-Tyr | D-Tyr | L-Tyr | D-Tyr | L-Tyr | D-Tyr | binding<sup>a</sup> | efficiency |
| Experimental variable | $K_M(\text{L})$ | $K_M(\text{D})$ | $k_{\text{cat}}(\text{L})$ | $k_{\text{cat}}(\text{D})$ | $\frac{k_{\text{cat}}}{K_M}(\text{L})$ | $\frac{k_{\text{cat}}}{K_M}(\text{D})$ | $\frac{K_M(\text{D})}{K_M(\text{L})}$ | $\frac{k_{\text{cat}}(\text{D})}{K_M(\text{D})} / \frac{k_{\text{cat}}(\text{L})}{K_M(\text{L})}$ |
| Reference state | apo enzyme |  | L-Tyr complex | D-Tyr complex | apo enzyme |  | L-Tyr complex | activated L complex |
| Target state | L-Tyr complex | D-Tyr complex | activated complex |  | activated complex |  | D-Tyr complex | activated D complex |
| Properties filtered | stability |  | stability |  | stability |  | stability, D-Tyr binding | stability, D-Tyr efficiency |

<sup>a</sup>Design for specific D-Tyr binding was not performed here.

We summarize the results in Table 5 and Fig. 4. We refer to the TyrRS variants by the identities of the four designed positions. Thus, the wildtype (WT) variant is DYQQ. Relative rates are characterized by 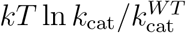. Catalytic efficiencies for the substrate S (L- or D-Tyr) are characterized by the quantity

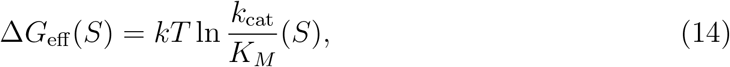

which can be interpreted as the free energy to bind and activate the substrate S.^41,54^ Specificities are characterized by the free energy difference Δ*G*_eff_ (*S*) − Δ*G*_eff_ (*S′*), where *S* and *S′* are the target and reference ligands, respectively. Table 5 shows results for four (out of 8) design criteria; for each one, the top four sequences are shown and values in the corresponding column are in boldface. Results with other criteria are in Supplementary Material, including design for a low reaction rate (transition state binding).

**Table 5:**
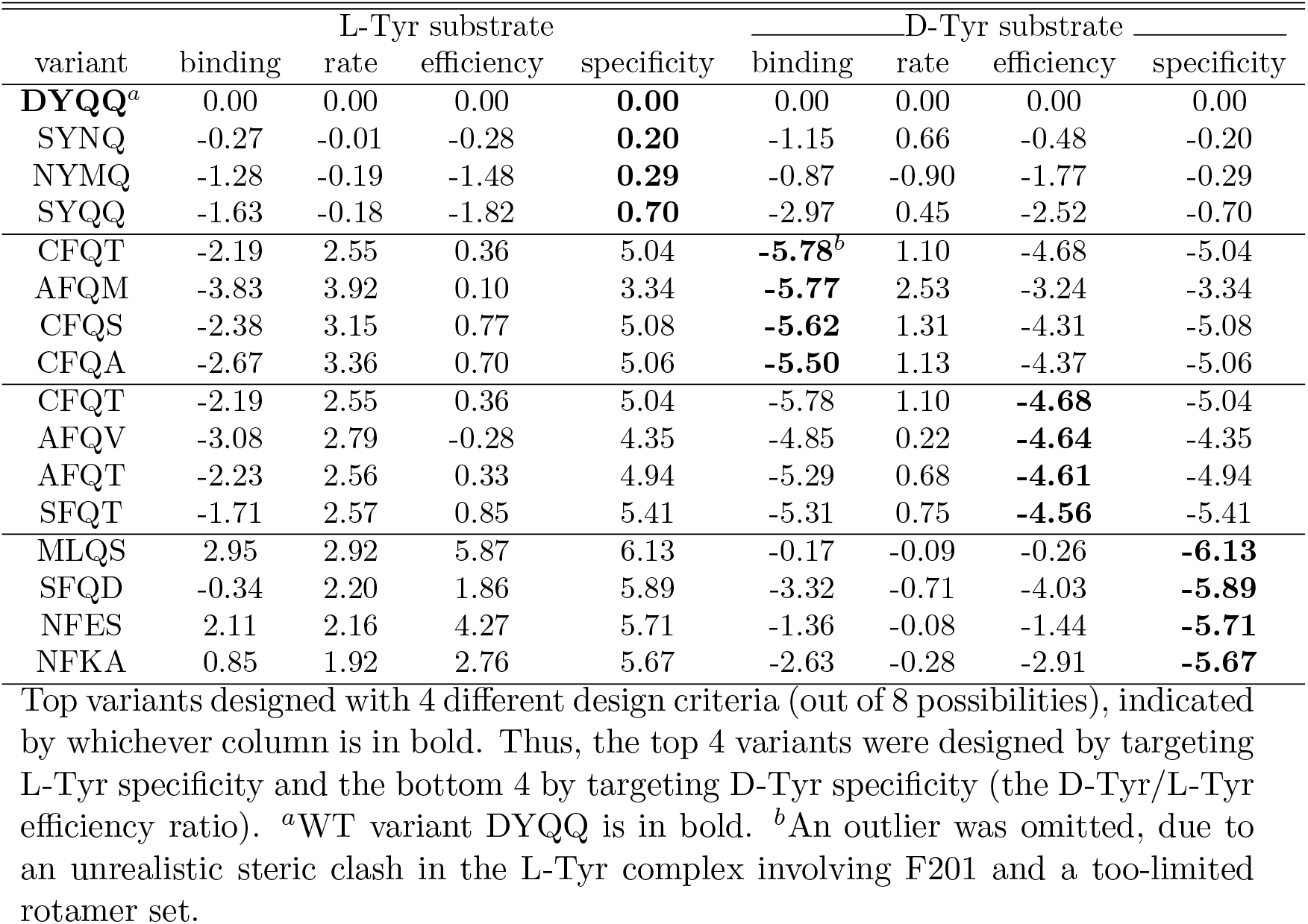
Top TyrRS designs with selected design criteria.

| variant | L-Tyr substrate |  |  |  | D-Tyr substrate |  |  |  |
| --- | --- | --- | --- | --- | --- | --- | --- | --- |
|  | binding | rate | efficiency | specificity | binding | rate | efficiency | specificity |
| **DYQQ<sup>a</sup>** | 0.00 | 0.00 | 0.00 | **0.00** | 0.00 | 0.00 | 0.00 | 0.00 |
| SYNQ | -0.27 | -0.01 | -0.28 | **0.20** | -1.15 | 0.66 | -0.48 | -0.20 |
| NYMQ | -1.28 | -0.19 | -1.48 | **0.29** | -0.87 | -0.90 | -1.77 | -0.29 |
| SYQQ | -1.63 | -0.18 | -1.82 | **0.70** | -2.97 | 0.45 | -2.52 | -0.70 |
| CFQT | -2.19 | 2.55 | 0.36 | 5.04 | **-5.78<sup>b</sup>** | 1.10 | -4.68 | -5.04 |
| AFQM | -3.83 | 3.92 | 0.10 | 3.34 | **-5.77** | 2.53 | -3.24 | -3.34 |
| CFQS | -2.38 | 3.15 | 0.77 | 5.08 | **-5.62** | 1.31 | -4.31 | -5.08 |
| CFQA | -2.67 | 3.36 | 0.70 | 5.06 | **-5.50** | 1.13 | -4.37 | -5.06 |
| CFQT | -2.19 | 2.55 | 0.36 | 5.04 | -5.78 | 1.10 | **-4.68** | -5.04 |
| AFQV | -3.08 | 2.79 | -0.28 | 4.35 | -4.85 | 0.22 | **-4.64** | -4.35 |
| AFQT | -2.23 | 2.56 | 0.33 | 4.94 | -5.29 | 0.68 | **-4.61** | -4.94 |
| SFQT | -1.71 | 2.57 | 0.85 | 5.41 | -5.31 | 0.75 | **-4.56** | -5.41 |
| MLQS | 2.95 | 2.92 | 5.87 | 6.13 | -0.17 | -0.09 | -0.26 | **-6.13** |
| SFQD | -0.34 | 2.20 | 1.86 | 5.89 | -3.32 | -0.71 | -4.03 | **-5.89** |
| NFES | 2.11 | 2.16 | 4.27 | 5.71 | -1.36 | -0.08 | -1.44 | **-5.71** |
| NFKA | 0.85 | 1.92 | 2.76 | 5.67 | -2.63 | -0.28 | -2.91 | **-5.67** |

Top variants designed with 4 different design criteria (out of 8 possibilities), indicated by whichever column is in bold. Thus, the top 4 variants were designed by targeting L-Tyr specificity and the bottom 4 by targeting D-Tyr specificity (the D-Tyr/L-Tyr efficiency ratio). <sup>a</sup>WT variant DYQQ is in bold. <sup>b</sup>An outlier was omitted, due to an unrealistic steric clash in the L-Tyr complex involving F201 and a too-limited rotamer set.

**Figure 4:**
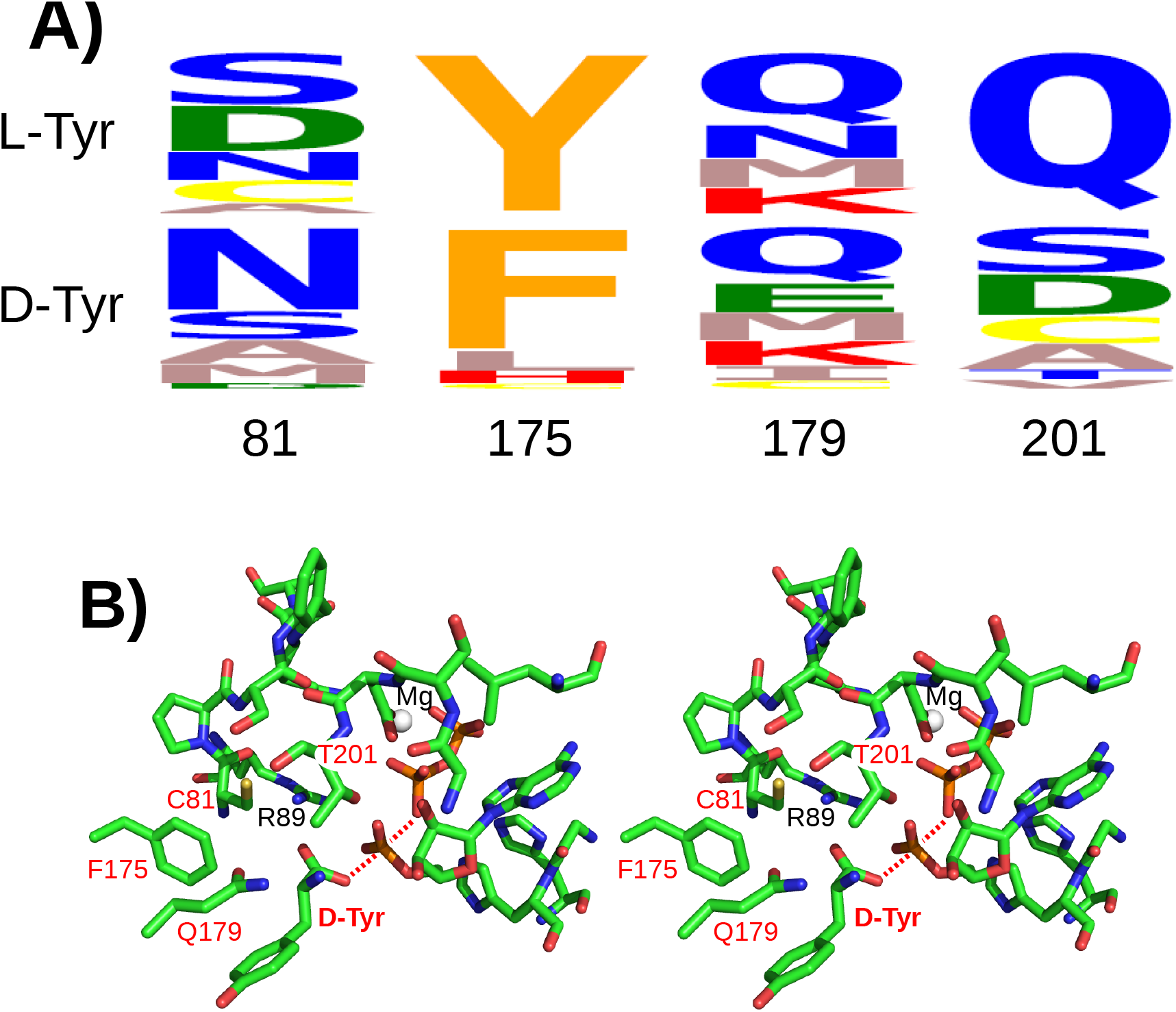
**A)** Sequence logos for TyrRS designs targeting specificity for L-Tyr (above) orD-Tyr (below). Specificity was defined by the ratio of catalytic efficiencies. **B)** Stereo view of the transition state complex of the CFQT variant, which maximizes D-Tyr binding and efficiency. Mutated residues have red labels. Mg and Arg89 are also labeled.

The upper section of Table 5 lists the top sequences obtained with specificity for L-Tyr as the design criterion. The WT sequence was ranked first for this property, which is satisfying. Some other variants yielded better values for other properties; thus, SYQQ had a better predicted L-Tyr affinity and efficiency and a slightly better reaction rate. Considering D-Tyr activity (catalytic efficiency), the top variant was CFQT, which was also the top D-Tyr binder (excluding an outlier), and had a predicted D-Tyr efficiency that was improved over WT (taken as zero) by 4.68 kcal/mol. All the top efficiency variants replaced Tyr175 by Phe, preserved the WT Gln175, and replaced Asp81 by a smaller, neutral side chain (C, A, or S). The top specificity variants were similar, with somewhat larger side chains at position 81. These included Asn, which has the same shape as Asp. These Asp81 changes to a neutral side chain appear consistent with the inverted specificity of the Asp81Arg single mutant discovered earlier. ^10^ The present predictions gave larger, more favorable changes in D-Tyr binding, efficiency, and specificity, presumably because the designs targeted these properties directly.

Sequence logos (Fig. 4A) show the amino acids preferred at each mutating position according to the specificity of the catalytic efficiency: preference for Lover D-Tyr (above) or the reverse (below). The WT types were most or second-most preferred for L-Tyr specificity. For D-Tyr specificity, Tyr175 was changed to Phe, Gln179 remained the preferred type, Asn81 replaced Asp as the preferred type, while S, D, C, A were preferred at 201. The 3D structure of the CFQT variant is shown in Fig. 4B. This variant maximized D-Tyr binding and efficiency. The Cys81 side chain occupies space freed up by the L-Tyr → D-Tyr change (which shifts the ligand ammonium to the right). Phe175 is in the same place as the WT Tyr, while Thr201 (replacing Gln) interacts nicely with the D-Tyr ammonium. Thus, the predicted structure appears plausible. Notice that for the WT L-Tyr complex, Proteus predicted side chain positions in the active site that superimposed closely on the X-ray conformation (not shown).

## 6 Discussion and perspectives

We have defined “physics-based” CPD or *ϕ*-CPD by its energy function and sampling strategy. With Proteus, we have primarily used the Amber ff99SB force field for proteins and ligands, a GB variant parameterized for this force field,^26^ and an SA or LK nonpolar term. LK was parameterized against structural and thermodynamic measurements, without any CPD data.^18^ There are two SA parameterizations: one based on the same data^18^ and one tuned for whole protein design.^47^ For whole protein design, an empirical unfolded model is used, as usual in CPD. The GBSA folded model and empirical unfolded model recently allowed us to successfully redesign a PDZ domain.^19^ For the design of ligand binding, the unfolded model is a non-empirical, extended peptide model. Proteus files (psf, force field parameters) are compatible with the NAMD program, so that designed sequences are readily studied with molecular dynamics simulations in explicit solvent.

Proteus uses REMC with long trajectories and Boltzmann sampling. This includes multibackbone situations where a new hybrid MC was specifically designed for Boltzmann sampling. ^38^ As a result, we can use importance sampling to target specific properties like binding and to estimate relevant free energies. Thus, constant-pH simulations yield pK_*a*_’s^17,55^ and constant-activity simulations yield binding affinities.^56^ Adaptive Wang-Landau MC is even more versatile, and was used to obtain conformational free energies, ^38^ acid/base constants,^57^ relative binding constants, catalytic rates, and catalytic efficiencies.^41^ Many capabilities were illustrated by the TyrRS design, above.

Calculations are efficient, thanks to the fixed backbone and discrete rotamers, and the energy matrix lookup table. The FDB method retains the many-body GB property, using a lookup table of lookup tables, with a cost about 5 times more than the usual pairwise additive GB simplification. Design for ligand binding (using the more accurate FDB), including system setup, energy matrix calculation, two adaptive MC simulations, and postprocessing is about a 24 hour operation on a desktop computer with 16 Intel cores. The adaptive bias works well with mutation spaces of 4-5 mutating positions and a few hundred thousand allowed sequences, especially if two-position bias terms are included. To explore more positions, we have used a strategy where we first explore pairs of positions involving 20 residues in the binding site (190 pairs). Then, positions that produced high scoring designs (as part of one or more pairs) are combined into sets of 4-5 positions; these are redesigned all at once. This two-tier approach can handle 20 positions within a pocket in a day if the simpler NEA GB is used for the first step (screening of pairs).

Some other functionalities were not described above but are documented elsewhere. ^16^ Proteus can reformat its own energy matrix for the optimization program Toulbar2, which does sequence optimization using sophisticated exact^58–60^ or heuristic methods.^61^ Proteus also implements its own rather effective, multiple minimization, heuristic method.^29,35^ Going in another direction, increased physical realism is provided by a method for design with a fully flexible backbone and continuous rotamers, which uses a hybrid MC/MD approach,^22^ although the current implementation is still slow.

Since *ϕ*-CPD uses a molecular mechanics energy, it is readily extended to unnatural amino acids, small nucleotides, and other small molecules. Importantly, this includes enzyme substrates in their activated, transition state. It can in principle be applied to RNA, but a fixed-backbone, discrete rotamer description is not yet available; work in this direction is underway. *ϕ*-CPD lends itself naturally to cycles that go back and forth between the CPD level of theory (fixed backbone, rotamers, implicit solvent) and fully flexible MD with the same force field but explicit solvent. Finally, *ϕ*-CPD can benefit from advances in molecular simulations and force fields and can yield physical insights and understanding. It represents a complementary route, alongside more empirical ones.

## Acknowledgements

We are grateful to several colleagues for helpful discussions and/or contributions to Proteus development: Théo Adrien, David Allouche, Edouard Audit, Sophie Barbe, Julien Bigot, Xingyu Chen, Juan Cortes, Marie-Pierre Dreanic, Nicolas Panel, Thomas Schiex, Seydou Traoré. We thank Axel Brünger for sharing the X-PLOR command parser.

## Supplementary material

Additional data are provided as a single Supplementary Material file, which describes in more detail the computational protocols used for the whole protein and TyrRS design applications and provides additional results.

## Table of contents graphic

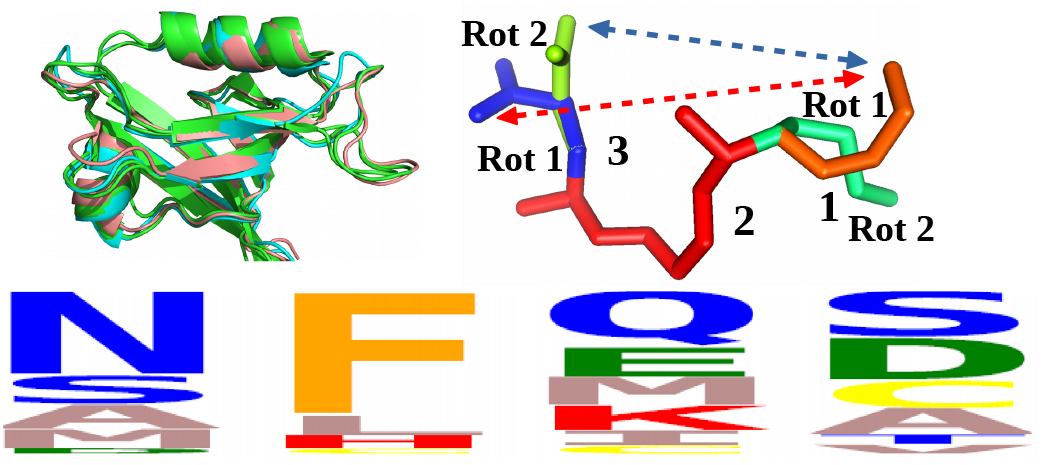

## Supplementary Material

## 1 Computational methods

Here we report details of the computational protocols used for the present applications.

### 1.1 Whole protein design

#### 1.1.1 Experimental amino acid frequencies

We considered 3 protein domains from the PDZ family (Table S1). To define the target amino acid frequencies, we collected homologous sequences for each domain by searching the Non-redundant (NR) database with NCBI/Blast [1], Blosum62 scoring matrix, and the PDB sequence as query. We retained homologues with sequence identities vs. the query above 60%. We used the HMMER algorithm [2] and the Superfamily tool [3] to identify and eliminate any Blast hits that did not belong to the same protein family as the query, leaving a total of 199 homologues. Finally, we aligned each query and its homologues with Clustal Omega [4].

**Table S1:**
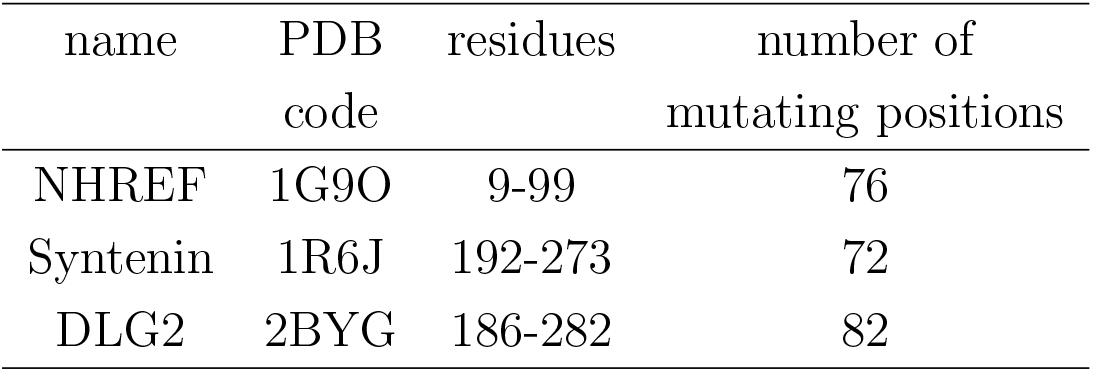
PDZ proteins targeted for redesign.

#### 1.1.2 Structure preparation and energy matrices

We began with the X-ray structures, listed in Table S1, and used the Modeler program to build any missing heavy atoms. We then placed hydrogens with the protX module of Proteus and did 1000 steps of conjugate gradient minimization with harmonic restraints of 3 kcal/mol/Å^2^ on heavy atoms. With the resulting backbone structure and side chain rotamers, we calculated the interaction energy matrix (IEM) with protX as previously [5]. For GBLK, we used parameters optimized earlier [6]: a protein dielectric constant *ϵ_P_* of 6.8, *λ* = 5.33, *S*^alk^ = 0.474, 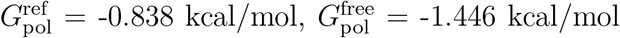. For GBSA, we used *ϵ_P_* = 8 or 4, and surface area parameters optmized earlier for CPD [7]. For the GB term, we used the Fluctuating Dielectric Boundary scheme (FDB) [8]. The IEM contains interactions between each residue and the backbone and between residue pairs (I, J), for all rotamers and for all side chain types except Gly and Pro. We used a slightly enhanced version of the Tuffery rotamer library [7, 9], along with “native” rotamers. Before the energy computation, for each state of residue I and residue pair I, J we performed 15 steps of minimization in order to alleviate bad steric contacts due to the rotamer approximation.

#### 1.1.3 *E*^uf^ optimization

The unfolded state energies were optimized by likelihood maximization, as previously [5]. Parameters were optimized separately for each protein family. Briefly, separate parameters were assigned to buried and exposed positions in the folded structures of the proteins. Rounds of REMC were done, where each protein was simulated twice, with either half of its positions allowed to mutate. After each round, MC amino acid frequencies from a low temperature replica (thermal energy of 0.26 kcal/mol) were compared to the experimental ones. Unfolded energies were updated with a linear update rule and a new round performed. To limit the number of adjustable parameters, early rounds grouped amino types into 11 classes, with shared parameters. In later rounds, each amino acid type had its own parameter.

#### 1.1.4 Sequence design and analysis

Sequence design was performed using the optimized reference energy set obtained for each protein family. All positions (except Gly, Pro) could mutate freely into all types except Gly, Pro. We did REMC with eight replicas for 750 million MC steps. MC moves included rotamer changes and mutations at one or two positions [5]. Thermal energies ranged from 0.175 to 3 kcal/mol. Periodic swaps were attempted between the conformations of two replicas *i, j* (adjacent in temperature) every 7.5 million MC steps. The 10,000 lowest energy sequences sampled by any replica were kept for analysis. Amino acid frequencies from MC for the PDZ proteins are compared to the experimental ones in Table S2. The converged reference energies are also given. The values with GBSA are in Table S3.

The sequences were submitted to the Superfamily library of hidden Markov models [3], which attempts to classify sequences according to the SCOP structural classification of proteins [10]. We also compared the designed sequences to natural sequences from the Pfam the “RP55” alignment for each family, using the Blosum40 scoring matrix. We first aligned the designed sequences to the Pfam alignment for each family, using the MAFFT program [11] and the Blosum40 scoring matrix.

The diversity in the 10000 designed sequences from each PDZ domain was computed using the position-dependent sequence entropy [5]:

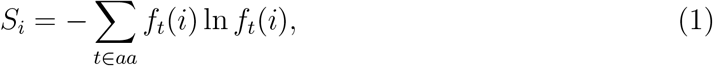

where *f_t_*(*i*) is the frequency of residue “type” *t* at position *i* of the designed sequences. Instead of the usual, 20 amino acid types, we employed six residue classes, corresponding to the following groups: {LVIMC}, {FYW}, {G}, {ASTP}, {EDNQ}, and {KRH}. The entropies are exponentiated, then averaged over all positions. To obtain the diversity of all 30000 designed sequences from the three domains, we initially organized them into a multiple sequence alignment.

### 1.2 Tyrosyl-tRNA synthetase design

#### 1.2.1 Transition state model

A model of the enzyme reaction transition state was prepared with the method described recently for methionyl-tRNA synthetase [12]. Briefly, we considered the enzyme reaction that transforms Tyr + ATP into tyrosyl adenylate (TyrAMP) and pyrophospate:

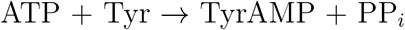

We assumed a transition state with pentavalent phosphorus coordination, including a Tyr backbone oxygen, with a trigonal bipyramidal geometry (Fig. S1. The axial PO equilibrium distance was set to 2.4 Å, based on *ab initio* optimization of a small fragment. RESP charges were computed at the HF/6-31G* level, as usual with Amber [13]. Other atom types and parameters were assigned by analogy to existing types in the force field.

**Table S2:**
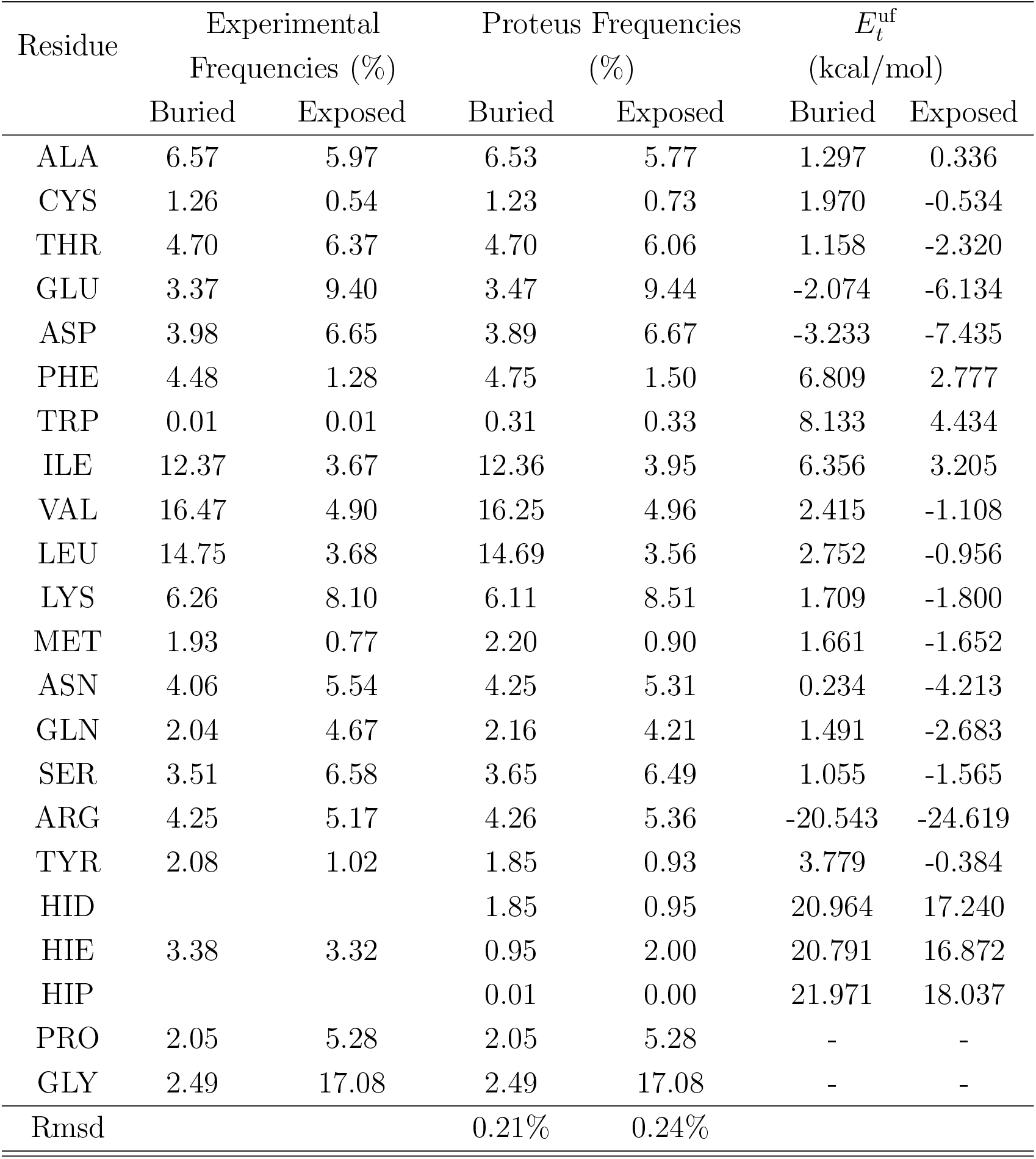
Amino acid composition of natural and designed PDZ sequences with GBLK.

**Table S3:** Amino acid composition of natural and designed PDZ sequences with GBSA(*ϵ_P_* =8)

| Residue | Experimental frequencies (%) | | Proteus frequencies (%) | | $E_t^{\text{uf}}$ (kcal/mol) | |
| --- | --- | --- | --- | --- | --- | --- |
|  | Buried | Exposed | Buried | Exposed | Buried | Exposed |
| ALA | 6.57 | 5.97 | 6.95 | 7.37 | 0 | 0 |
| CYS | 1.26 | 0.54 | 1.31 | 2.04 | -2.137 | -2.113 |
| THR | 4.70 | 6.37 | 4.77 | 7.47 | -5.736 | -8.712 |
| GLU | 3.37 | 9.40 | 3.65 | 10.64 | -15.956 | -20.69 |
| ASP | 3.98 | 6.65 | 4.24 | 7.95 | -16.115 | 20.17 |
| PHE | 4.48 | 1.28 | 4.45 | 2.54 | 3.142 | 1.865 |
| TRP | 0.01 | 0.01 | 0.35 | 1.41 | 0.750 | -2.532 |
| ILE | 12.37 | 3.67 | 12.05 | 3.01 | 4.622 | 1.472 |
| VAL | 16.47 | 4.90 | 16.59 | 5.90 | 0.523 | -2.455 |
| LEU | 14.75 | 3.68 | 15.16 | 5.05 | -0.243 | -4.277 |
| LYS | 6.26 | 8.10 | 7.01 | 9.56 | -4.966 | -8.418 |
| MET | 1.93 | 0.77 | 2.07 | 2.35 | -1.918 | -4.010 |
| ASN | 4.06 | 5.54 | 4.44 | 6.94 | -15.654 | -20.31 |
| GLN | 2.04 | 4.67 | 2.38 | 5.81 | -15.263 | -20.556 |
| SER | 3.51 | 6.58 | 3.72 | 7.95 | -6.392 | -8.063 |
| ARG | 4.25 | 5.17 | 4.76 | 6.72 | -45.249 | -49.887 |
| TYR | 2.08 | 1.02 | 2.45 | 2.52 | -6.385 | -9.791 |
| HID |  |  | 1.66 | 2.70 | 10.055 | 11.488 |
| HIE | 3.38 | 3.32 | 1.96 | 0.20 | 9.887 | 5.551 |
| HIP |  |  | 0.0 | 0.0 | 11.370 | 6.746 |
| PRO | 2.05 | 5.28 | 2.05 | 5.28 | - | - |
| GLY | 2.49 | 17.08 | 2.49 | 17.08 | - | - |
| Rmsd |  |  | 0.23% | 0.87% |  |  |

**Figure S1:**
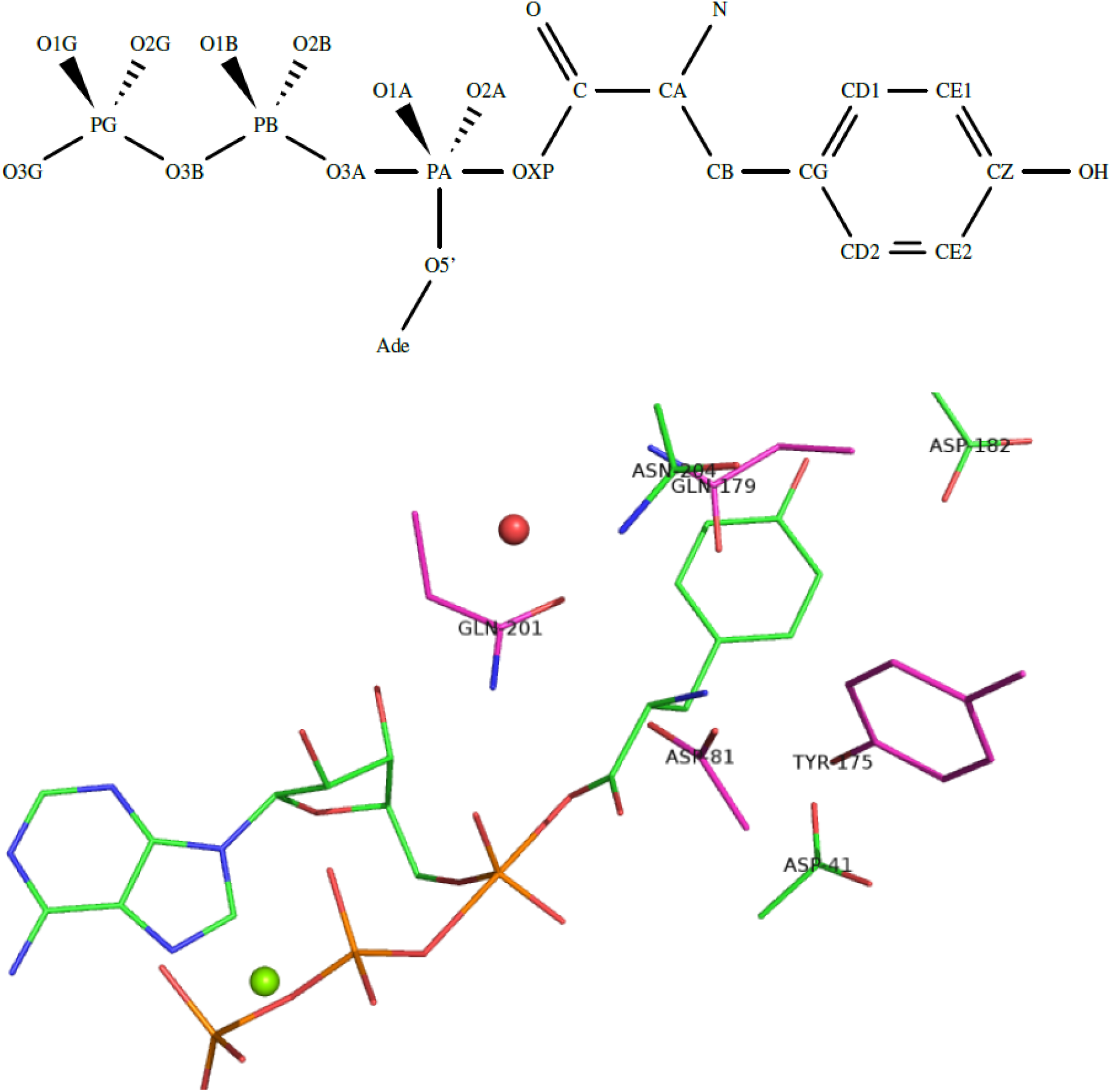
Above: the transition state ligand. Below: Closeup of the activated complex. A buried water and an ATP-bound magnesium are shown as spheres. Four mutating positions are labeled, as well as nearby Asp41 and Asp182.

#### 1.2.2 Structure preparation

Structural models of the TyrRS protein were prepared, alone and in complex with L- or D-Tyr or transition states. The TyrRS model was based on the 1VBM PDB entry, an *E. coli* TyrRS structure in complex with an L-TyrAMP analog. Only the A chain was used, stripped of water molecules, ions, and ligands, except one conserved, buried water molecule [14]. The four ligands considered were the reactants of the adenylation reaction, that is ATP plus either L- or D-Tyr, as well as the L and D transition states, denoted LTP and DTP. Ligand conformations were prepared, starting from the L-TyrAMP analogue present in the 1VBM structure. The analogue was first converted to a real L-TyrAMP by modifying the appropriate atoms. L-Tyr and AMP molecules were separated. ATP was then built by combining the AMP extracted from 1VBM with the second and third phosphates extracted from the ATP molecule present in the 1H3E PDB entry, after superposition on the AMP. 1H3E is a *Th. thermophilus* TyrRS structure in complex with tRNA, ATP, and tyrosinol. Finally, the LTP structure was built by combining ATP and L-Tyr, and modifying the first phosphate (PA, O1A, O2A, O5’ atoms) geometry to make it trigonal bipyramidal (PA, O1A, O2A, O5’ in the same plane; OXP-PA-O3A axis perpendicular to the plane). The distance of PA to OXP and O3A was 2.2 Å, while the distance of PA to O1A, O2A, and O5’ was 1.5 Å, close to the equilibrium values. The D-Tyr version of the ligands was prepared by rotating the N atom by 120^*◦*^ around the CA-C axis. A magnesium ion was included in all ligand models, positioned between the second and third phosphates. Protonation states were standard, except for Asp182, close to the substrate side chain, which was protonated, based on previous PropKa calculations [15] and structure inspection [14]. The models were built and minimized with the protX module of Proteus.

#### 1.2.3 Energy matrix calculation

An interaction energy matrix was calculated for each model, using protX. We followed the same protocol as in previous work [7, 16], where details are given. Briefly, the rotamer library was prepared from the 1995 library of Tuffery et al [9], the energy function was of the GBSA form, the molecular mechanics force field was AMBER ff99SB for the protein [17, 18], and compatible parametrs were used for ATP [19] and Tyr [20]. For the transition state, see above. A GB/HCT variant was used [21] with a protein dielectric constant of 8 and the NEA procedure. Surface energy coefficients were previously optimized for CPD [7]. For the ligands, we constructed rotamers by grafting the usual Tyr side chain rotamers onto each ligand backbone and keeping the rest (backbone, AMP moiety) fixed. With other, nonpeptidic ligands, “rotamers” can be provided in the form of a collection of poses, grouped in a PDB file.

#### 1.2.4 Sequence design

We redesigned four amino acids close to the Tyr ligand backbone, as previously [14, 22]: Asp81, Tyr175, Gln179 and Gln201. Mutations into all types were allowed except Y and W at position 201, due to a lack of space in the binding pocket. The other positions kept their native types. The conformation of amino acids whose side chain was within 8 Å of the ligand’s Tyr moiety was allowed to vary. Sequences and conformations were explored by MC with the protMC module of Proteus. Simulations were run for 10^9^ steps, with thermal energy 0.6 kcal/mol and move probabilities of 0.9, 0.1, 0.9, and 0.1, respectively for single rotamer, single mutation, pair of rotamers, and mutation-rotamer moves. An adaptive bias was optimized in certain simulations, with parameters reported earlier [12].

The adaptive landscape flattening method [12, 23] can be used in two ways to calculate a free energy difference between two states. In the first approach, a bias potential is obtained by an adaptive Wang-Landau procedure for one of the two states, called the reference state. The bias has the form of position-specific amino acid energies that flatten the free energy landscape in sequence space. The second state is then sampled with the bias potential applied, directly producing sequences populated according to the free energy difference between the two states. In the second, indirect approach, the free energy landscape of *both* states is flattened by developing a separate bias potential for each state. The two states, say A and B are then simulated, each with its own bias potential. For two sequences *S* and *R* sampled in both states, the relative free energy difference between the two states can then be deduced from the sampled populations:

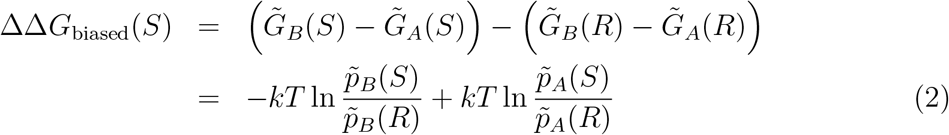

where *p* is the probability of observing sequence *S* or *R* in state A or B and the tilde sign indicates the bias is present. In this work, we followed the indirect approach, which gives more flexibility to analyze the results, such as the possibility of multi-criteria filtering.

## 2 Results

### 2.1 Tyrosyl-tRNA synthetase design

We provide some additional results in this section. Fig. S2 shows sequence logos obtained according to 8 design criteria: L- or D-Tyr binding, catalytic rate, efficiency, and specificity. With affinity design, a Tyr is found at position 175 for the L-Tyr and D-Tyr ligands, suggesting that this position is not able to discriminate them in terms of affinity. At the other mutating positions, the designs differs. Phe, Met, and Met are obtained for L-Tyr at positions 81, 179, and 201. For D-Tyr, we found Met, Val, and Phe at positions 81, 179, and 201. With catalytic rate design, with L-Tyr, we find Thr/Ser/Asp, Tyr, Ile/Met, and Gln at positions 81, 175, 179, and 201. For D-Tyr, we obtain Ser/Ala/Asn, Phe/Tyr, Met/Ile, and Asp/Cys at positions 81, 175, 179, and 201.

With catalytic efficiency design, with L-Tyr, we obtain Ser/Ala/Cys, Tyr, Gln/Met/Lys, and Gln/Ile at positions 81, 175, 179, and 201. The native types are thus retrieved for three positions (175, 179, and 201) out of four. This could indicate that these positions have been selected by evolution for catalytic efficiency. This is also an element in favor of the validity of our transition state model. For D-Tyr, we obtain Ala/Ser/Cys, Phe, Gln/Lys, and Thr/Cys/Ser at positions 81, 175, 179, and 201. We note that the profiles obtained for L- and D-Tyr are quite similar, suggesting that the effects contributing to catalytic efficiency are not strongly stereospecific. Finally, with specificity in favor of L-Tyr as the design target, the WT sequence is top-ranked (see main text), which is satisfying. With specificity in favor of D-Tyr, there are differences at positions 81, 175, and 201, as discussed in the main text.

**Figure S2:**
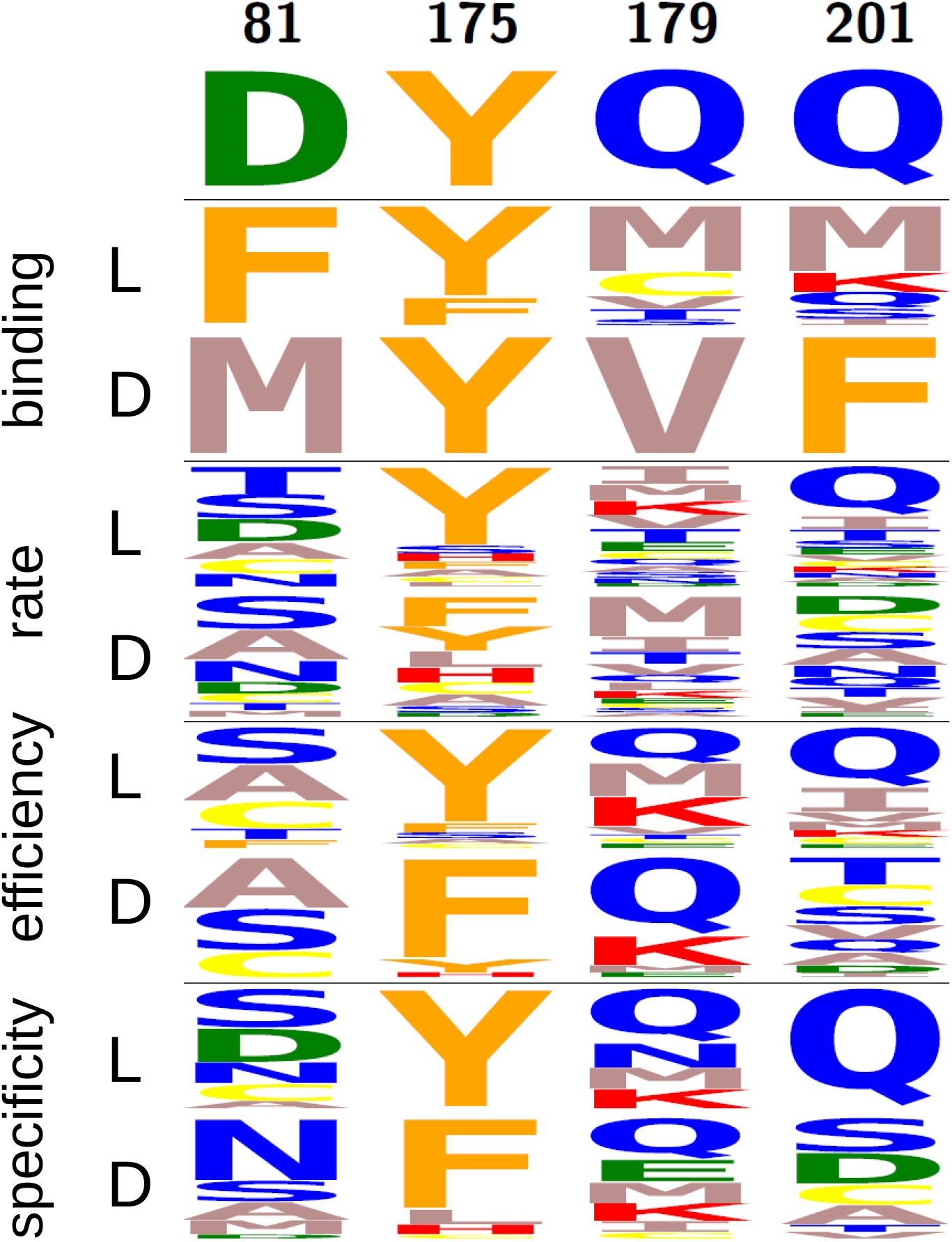
Sequence logos from TyrRS design with different criteria.

